# NK cells contribute to resistance to anti-PD1 therapy in immune-excluded melanomas

**DOI:** 10.1101/2023.12.14.571631

**Authors:** Ewout Landeloos, Joanna Pozniak, Niccoló Roda, Amber De Visscher, Asier Antoranz Martinez, Yannick Van Herck, Greet Bervoets, Francesca Bosisio, Veerle Boecxstaens, Ayse Bassez, Diether Lambrechts, Patrick Matthys, Oliver Bechter, Jean-Christophe Marine

**Affiliations:** Laboratory for Molecular Cancer Biology, Center for Cancer Biology, VIB, Leuven, Belgium; Laboratory for Molecular Cancer Biology, Department of Oncology, KULeuven, Leuven, Belgium; Department of General Medical Oncology UZ Leuven, Belgium; Department of Microbiology, Immunology and Transplantation, Rega Institute, Immunity and Inflammation Research Group, Immunobiology Unit, KULeuven, Leuven, Belgium; Laboratory of Translational Cell and Tissue Research, Department of Pathology, KULeuven, Leuven, Belgium; Department of Pathology, UZLeuven, Belgium; Department of Surgical Oncology, Oncological and Vascular Access Surgery, UZ Leuven, Leuven, Belgium; Laboratory of Translational Genetics, Center for Cancer Biology, VIB, Leuven, Belgium; Center for Human Genetics, KULeuven, Belgium

**Author notes:** These authors have contributed equally to this work (listed in alphabetical order) (Ewout Landeloos, Joanna Pozniak).

**Keywords:** NK cells, PD1 checkpoint immunotherapy, immune-excluded phenotype, melanoma

## Abstract

Immune checkpoint blockade (ICB) has become a standard of care in the treatment of metastatic melanoma (MM). Although ICB is particularly successful in some MM patients, more than half do not obtain a durable benefit. Biomarkers that predict response are urgently needed and overcoming intrinsic resistance is key to improving the success of ICB therapy. Using single cell RNA sequencing, we characterized the immune landscape of pre- and early on-treatment biopsies taken from a cohort of MM patients (n>20) exposed to ICB therapy. Our analysis identified >20 immune cell types and confirmed previously described associations between the abundance of various CD8 T cell populations and ICB outcome. Unexpectedly, we found that lack of response was associated with an increased occurrence of a granulysin-expressing (GNLY+) natural killer (NK) cell population. This observation was replicated in other MM cohorts and in a breast cancer cohort in which paired biopsies were also collected pre and early-on ICB therapy. Spatial proteomics revealed that whereas NK cells colocalized with CD8 T cells within the tumour bed in responding lesions, these cells accumulated at the tumour margin in non-responding lesions. Strikingly, depletion of NK cells in an NRAS-driven melanoma mouse model, which exhibits an immune-excluded phenotype and is refractory to ICB, promoted massive immune cell infiltration and tumour clearance upon anti-PD1 exposure. These data highlight a differential immune cell topography between early on-treatment responding and nonresponding MM lesions, which could be exploited to develop a robust stratification biomarker, and unravel an unexpected contribution of NK cells in primary resistance to ICB.

## Introduction

Immune checkpoint blockade (ICB) has significantly improved the survival of cancer patients and, in particular, of patients with metastatic melanoma (MM). Anti-PD-1 (aPD1) therapy, either alone or in combination with anti-CTLA-4 (aCTLA4), has become a standard of care for MM patients^1,2^. Despite remarkable clinical success, approximately half melanoma patients do not achieve disease control under ICB. Biomarkers that predict response to this treatment are urgently needed to spare non-responders from unnecessary toxicity and reduce health care costs. Moreover, understanding and targeting the mechanisms driving intrinsic resistance is key to improving the success of ICB^3^.

The success of ICB therapy largely depends on the cellular composition of the tumor immune microenvironment (TIME). Most, if not all, immune cells can dynamically transit between various cell states and thereby exhibit a range of phenotypes^4^ that can either promote or suppress antitumor immunity^5^. The efficacy of ICB therefore relies on the relative abundance between the immunosuppressive and immunostimulatory immune cell subsets. Bulk and, more recently, single-cell RNA sequencing (scRNA-seq) technologies allowed an in-depth dissection of the TIME and its evolution under ICB. This led for instance to the identification of specific CD8^+^ T cell subtypes that associate with ICB response, such as TCF7^+^ CD8^+^ T cells^6^ and EOMES^+^ CD69^+^ CD45RO^+^ effector memory T cells^7^. The presence, myeloid-derived suppressor cells (MDSCs) and tumour-associated macrophages (TAMs) has been linked to resistance^8,9^. In contrast, B lymphocytes, and natural killer (NK) cells have been associated with favourable response^10–14^. NK cells are considered important effectors in anti-tumour immune responses both in melanoma^15^ and in other cancer types, and have been shown to recruit stimulatory cDCs to the TIME^16–18^.

Unfortunately, however, the vast majority of the clinical studies aimed at understanding the contribution of various immune cells as mediators/modulators of the response to ICB in melanoma have thus far been conducted on samples collected from patients who had developed resistance to other therapeutic modalities, whereas ICB therapy is now used as a first line treatment. Moreover, these studies have often analysed biopsies collected before ICB administration and/or at disparate and late (i.e. at progression) on-treatment time points^19^. This is unfortunate because emerging evidence indicates that samples taken early during-treatment are likely to be more predictive than pre-treatment biopsies^20^. Finally, although the immune cell topography of solid tumours is increasingly recognized as an important predictive factor for response to immunotherapy^21,22^, most of the studies to date were performed on dissociated tumour samples and therefore, almost invariably, lack spatial context.

We therefore reasoned that a single-cell and spatially resolved dissection of the melanoma immune landscape of matching pre- and early on-treatment biopsies from drug-naïve MM patients undergoing ICB therapy may provide new critical insights into the mechanisms underlying intrinsic resistance to ICB and guide the development of effective combination therapies and/or robust stratification (composite) biomarkers.

## Results

### Dissecting the immune compartment of the melanoma ecosystem

To draft a temporally and spatially-resolved map of the immune landscape of melanoma under ICB, we prospectively collected paired MM biopsies (cutaneous, n=12; subcutaneous, n=8 and lymph node, n=26) before treatment (BT) and early (±2 weeks, before administration of the second ICB cycle) on-treatment (OT) from a cohort of 23 treatment-naïve (stage III-IV) patients (SPECIAL trial; UZ/KU Leuven #s62275)^23^. These patients receiving either anti-PD-1 monotherapy (nivolumab, n=17) or anti-PD-1 in combination with anti-CTLA-4 (ipilimumab + nivolumab, n=6). In total, we obtained matched biopsies across both time points for 20 out of 23 patients. Fresh tissue was processed for scRNA-seq, and fixed material was used for routine pathological assessment and targeted spatial proteomics (**Figure 1A**). Peripheral blood samples were collected from all patients and time points and used for immunophenotyping by flow cytometry (FACS).

**Figure 1:**
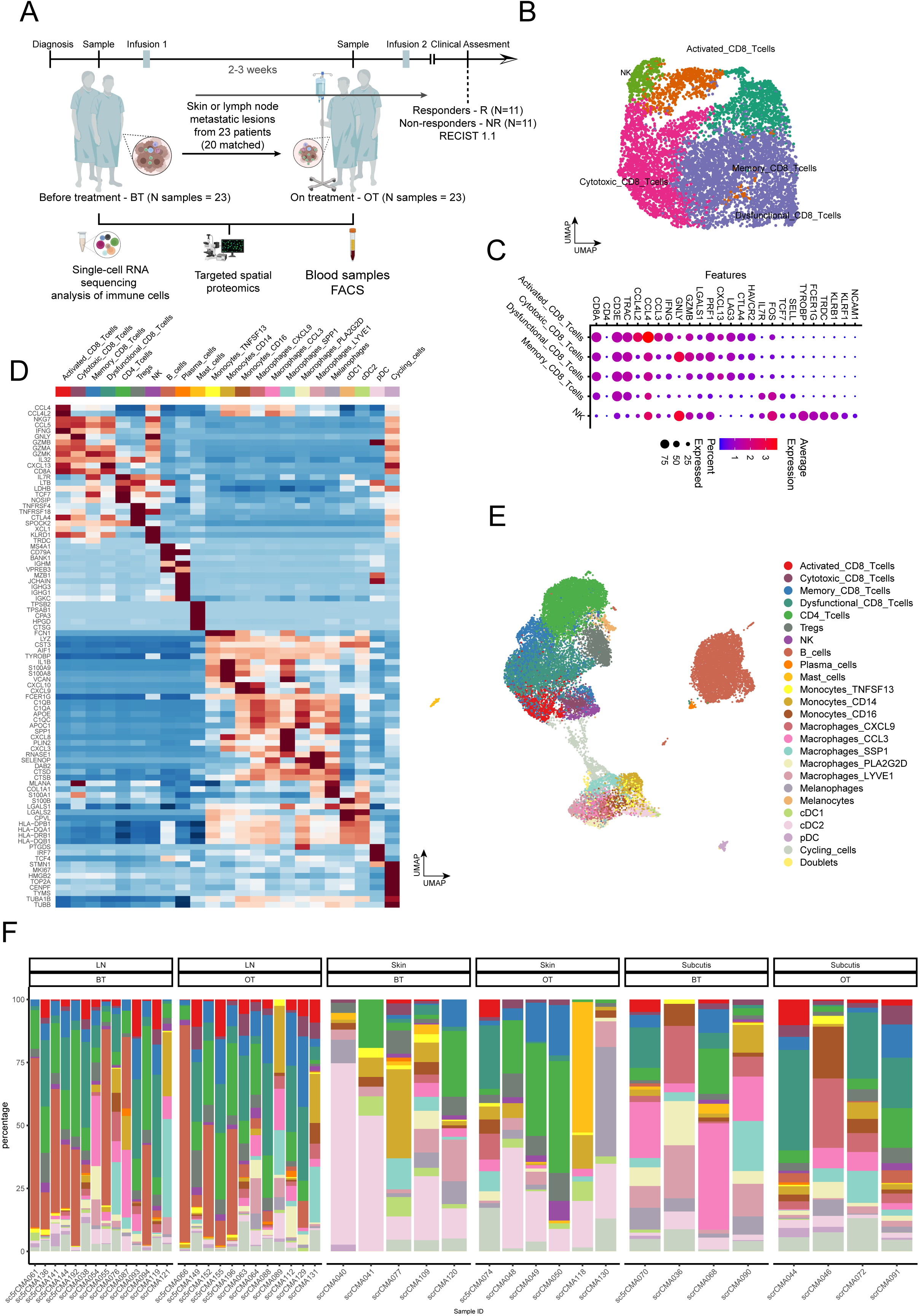
Dissecting the immune compartment of the melanoma ecosyste. **A,** Schematic describing the design of the SPECIAL clinical study, the timing of sample collection, number of patients enrolled and samples collected, and single-cell (spatial) methodologies deployed for their in-depth characterization **B,** UMAP of the CD8T states and NK cells identified by unsupervised Louvain clustering of the subset of CD8Tcells **C,** Dotplot of the top representative discriminatory marker genes of the cells presented in the panel B **D,** Heatmap representing the top 10 discriminatory marker genes of identified immune cells. **E,** UMAP of all immune cell types and states identified by unsupervised Louvain clustering. **F,** Proportions of the immune types of the total immune cell number presented in panel C in each sample grouped by metastatic location and timepoint (BT: Before Treatment, OT: On Treatment).

Patient clinical characteristics were obtained at baseline, and response (and progression) to ICB treatment was considered according to RECISTv1.1 criteria^24^. Patients with unresectable disease (n=18) achieving complete remission (CR) or partial remission (PR) as best overall response (BOR) were stratified as responders (Rs), whereas non-responders (NRs) achieved stable disease (SD) or progressive disease (PD). For the few patients with resectable disease (n=5), a first biopsy was obtained pre-treatment and a second one (surgical resection) was obtained after with 2 cycles of ICB. Presence or absence of complete pathological response (pCR; defined as the complete absence of residual viable tumour) was determined by analysis of the resection specimen (**Table S1A**).

After quality control, data normalization, integration and clustering, 25.401 single cells were identified as immune cells based on a high immune gene set activity^25^ and low mean copy number variant (CNV) score^26^. One sample was discarded because it contained less than 10 immune cells. Initial unsupervised Louvain clustering (harmony integration) identified 17 immune cell clusters, which were annotated based on previously published cell type specific gene signatures^6^ and expression of hallmark genes resulting in 10 main immune cell types. These included CD8+ and CD4+ T cells, Regulatory T cells, CD8+ T cells/NK cells, B cells, Plasma cells, Myeloid cells, Mast cells, and Plasmacytoid dendritic cells (pDC; **Figure S1A, B, Table S2**). We also identified a small population of cells expressing high levels of pigment production genes (*MLANA*, *PMEL*), and genes involved in extracellular vesicle formation (e.g., *CD63*, *VIM*, *TIMP3;* **Supplemental Table S1**). We annotated these cells as normal melanocytes because they were predominantly detected in biopsies originating from skin metastatic lesions and transcriptionally resembled normal melanocytes. Note that melanocytes do express several innate immune genes ^27^, which explain why they had not been filtered out.

Cells from the CD8+ T cell/NK cell and Myeloid cell clusters were further subclustered. In the CD8^+^ T cell/NK cell cluster 55% of cells were positive for *CD8A*, whereas 92% of cells expressed *GNLY*, suggesting that both cytotoxic T cells and cytotoxic NK cells populate this cluster (**Table S2**). We therefore re-clustered all CD8+ T cell clusters, thus retrieving a separate NK cell cluster characterized by high expression of *TYROBP*, *TRDC*, *NCAM1* (CD56) and low expression of *CD8A*, *CD4,* and *CD3E* (**Figure 1B, C** **and Table S3)**. Cells from this cluster displayed a cytotoxic phenotype, characterized by high expression of *GNLY*, *GZMB* and *PRF1*, ruling out overlap with other innate lymphoid cell populations^28^. In parallel, re-clustering identified four CD8+ T cells states: Memory (*IL7R*, *FOS*, *TCF7*, *SELL*)^5^, Activated (*IFNG, CCL3, CCL4*), Cytotoxic (*GNLY*, *GZMB*, *PRF1*), and Dysfunctional (expressing higher levels of *CXCL13, LAG3, CTLA4, and HAVCR2,* and lower levels of cytotoxic genes and *IFNG*) ^29^.

Similarly, since members of the mononuclear phagocyte system (MPS) display transcriptional similarities^30^, we re-clustered the Myeloid and pDC cluster together, resulting in 5 subclusters, corresponding to Macrophages_*C1Q*, Macrophages_*SPP1*, Monocytes, Dendritic cells, and pDCs respectively (**Figure S1C, Table S4**). However, Macrophage_*C1Q* cluster both contained cells expressing genes associated with an anti-inflammatory (*C1Q, APOE, MRC1*) and a pro-inflammatory role (*CLXL9,* HLA genes), leading us to further re-cluster this group of cells. We obtained five subclusters, named Macrophages_*LYVE1*, Macrophages_*CXCL9*, Macrophages_*PLAG2A*, Macrophages_*CCL3* and Melanophages (**Figure S1D; Table S5**). The Macrophages_*LYVE1* cluster largely overlapped with a known perivascular, tissue-resident, M2-like phenotype, expressing both vascular growth factors involved in angiogenesis (*CD209*, *LILRB5*, *PDGFC*) and genes associated with M2 polarization (*CD163*, *CSF1R*, *MRC1*) (**Table S5**). The Macrophages_*CXCL9* and Macrophages_*CCL3* gene signatures overlapped with the signature of M1-like macrophages. Note that, rather than being correlated with conventional M1 and M2 markers, the polarity of TAMs was shown to be associated with divergent expression in *CXCL9* and *CXCL10* versus *SPP1*^31^. The Macrophage_*CCL3* cluster was characterized by both pro-inflammatory cytokines (*CCL3*, *CCL4*, *CXCL3*, *TNF*) and transcription factors (*FOS*, *JUN*, *ATF3*, *NFKB* genes) associated with macrophage activation (**Table S5**)^32,33^.

Finally, because of their known phenotypic similarities, plasticity, and complex developmental trajectories, we re-clustered Monocytes and Dendritic cells together^34^. This analysis revealed clusters corresponding to classical monocytes (Monocytes_*CD14*), non-classical monocytes (Monocytes_*CD16*), intermediary monocytes (Monocytes_TNFSF13), conventional type 1 dendritic cells (cDC1) and conventional type 2 dendritic cells (cDC2). (**Figure S1E, Table S6**)^30,35^.

In summary, we identified 22 different immune cell types (**Figure 1D, E, Table S7**) within our MM biopsies, including four CD8+ T cell states, Tregs, CD4+ T cells, B cells, Mast cells, 3 Monocyte, 6 Macrophage and 3 Dendritic cell subclusters.

As expected, some cell clusters were overrepresented in certain tissues of origin **(Figure 1D, Figure S1F).** For example, B cells were almost exclusively detected in lymph node biopsies, whereas myeloid cells were more common in skin and subcutaneous metastases. This was consistent across time points (**Figure S1F).** Importantly, biopsies from different tissues were evenly distributed across treatment outcomes (Chi2 test p value = 0.7, **Figure S1G**).

### Cytotoxic NK cell abundance is associated with intrinsic resistance to ICB

We next tested whether the relative proportion of each specific immune cell type/subset correlated with clinical response (**Figure 2A, Figure S2A**). Such an association was only detected for two distinct immune cell types in this dataset.

**Figure 2:**
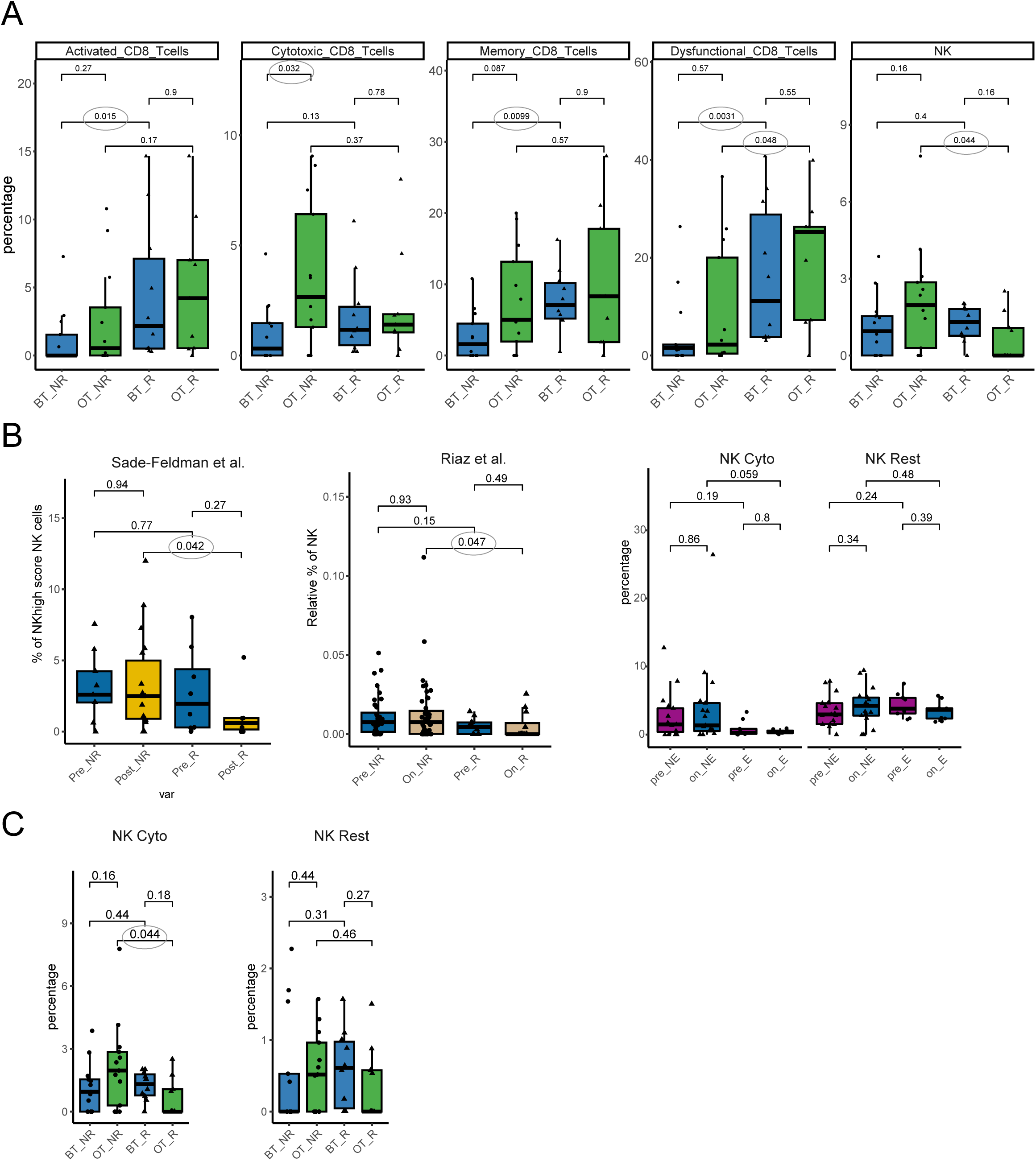
Cytotoxic NK cell abundance is associated with ICB resistance. **A,** Percentage of CD8 T-cell subsets and NK cells out of all immune cells compared among before treatment (BT) and on treatment (OT) samples and both response groups (two-sided Wilcoxon test, circle = p<0.05). **B,** Validation of NK cells association with lack of response in single cell RNA-seq melanoma cohort^6^ identified based on NK cell score acquired from our data (left), in bulk RNA-seq melanoma cohort^37^ identified based on CIBERSORTx (middle), and in the single cell RNA-seq breast cancer cohort identified as in the original study (right)^39^. **C,** Percentage of NK_cyto_ and NK_rest_ score measured in NK cells out of all immune cells and compared among before treatment (BT) and on treatment (OT) samples and both response groups (two-sided Wilcoxon test, circle = p<0.05).

Consistent with previous studies^7,36^, we found that CD8+ T cells exhibiting an Activated, Memory, and Dysfunctional profile were more abundant in lesions from Rs than in NRs (**Figure 2A**). These differences were less pronounced at the on-treatment (OT) timepoint. Notably, we observed an increase in Cytotoxic CD8^+^ T cells in NR OT, while the amount of Cytotoxic CD8+ T cells remained stable in R patients over the course of treatment. Unexpectedly, NK cell abundance OT was significantly higher in lesions from NRs compared to Rs (**Figure 2A**). Strikingly, this observation could be recapitulated in other publicly available datasets. The first dataset we mined was a scRNA-seq dataset containing 16.291 individual immune cells from 48 biopsies collected pre- and post-treatment (wide range of timepoints between days 22-867) from MM patients exposed to aPD-1 therapy^6^. Consistent with data from our cohort, a significant increase in the percentage of NK cells was observed in OT lesions from NRs compared Rs (**Figure 2B, left**). Additionally, we leveraged a large bulk RNA-seq dataset from a cohort of MM patients treated with aPD1 therapy^37^, and deconvolved the data using CIBERSORTx^38^ to estimate the percentages of NK cells in each sample. Sampling in this dataset was done pre-treatment and on-treatment (between days 23-29). Again, the NK cell percentage OT was significantly higher in NRs compared to Rs (**Figure 2B, middle).** These data indicated that a relatively high abundance of NK cells in (early) on-treatment biopsies predicts lack of response to ICB in patients with MM. Importantly, this finding may be generalizable as it could be replicated in other cancer type. We indeed observed a similar association in a scRNA-seq dataset of breast cancer patients treated with aPD1 (pembrolizumab) and for which samples were processed similarly and at the exact same time points, namely pre-treatment and before the second ICB infusion^39^. In this study, the authors identified two populations of NK cells, termed NK_CYTO_ and NK_REST_. Strikingly, the percentage of NK_CYTO_ was associated with a trend towards limited or no clonotype T cell expansion both BT and OT **(Figure 2B, right)**. Such association was absent for the NK_REST_ cell population. We next scored the NK_CYTO_ and NK_REST_ signatures from the breast cancer dataset within the NK cell population from our melanoma dataset. Interestingly, the percentage of cells with a positive score for the NK_CYTO_ signature was higher in OT MM lesions from NRs. In contrast and similarly to what we observed in the breast cancer cohort, no association with response was found for the cell expressing the NK_REST_ signature (**Figure 2C**).

### Peripheral blood NK cells display a differently activated phenotype in Rs vs NRs

We next investigated whether ICB therapy was also associated with changes in the pool of peripheral blood NK cells (PBNKs). We performed multiparametric flow cytometry using the blood samples from our cohort: 28 NR samples (BT, n = 13; OT, n = 15) and 23 R samples (BT, n = 12 OT, n = 11). NK cells were defined as CD19- CD14- CD123- CD3- CD56+ cells **(Figure S3).** We found no difference in the percentage of total NK cells, nor in CD56^Bright^ and CD56^Dim^ subsets, across time points and ICB outcomes^40^ (**Figure 3A**). Notably, although the ICB treatment did not induce a significant upregulation of tumour necrosis factor (TNF)–related apoptosis-inducing ligand (TRAIL) in the blood Ns nor NRs patients, we observed that the peripheral fraction of Granzyme B+ and Perforin+ NK cells decreased in NR (but not in R) patients during treatment (**Figure 3B**). These data may be consistent with an increased recruitment from the blood to the TIME of Granzyme B+ and Perforin+ NK cells in NRs than in Rs, possibly explaining their increased abundance in NR lesions.

**Figure 3:**
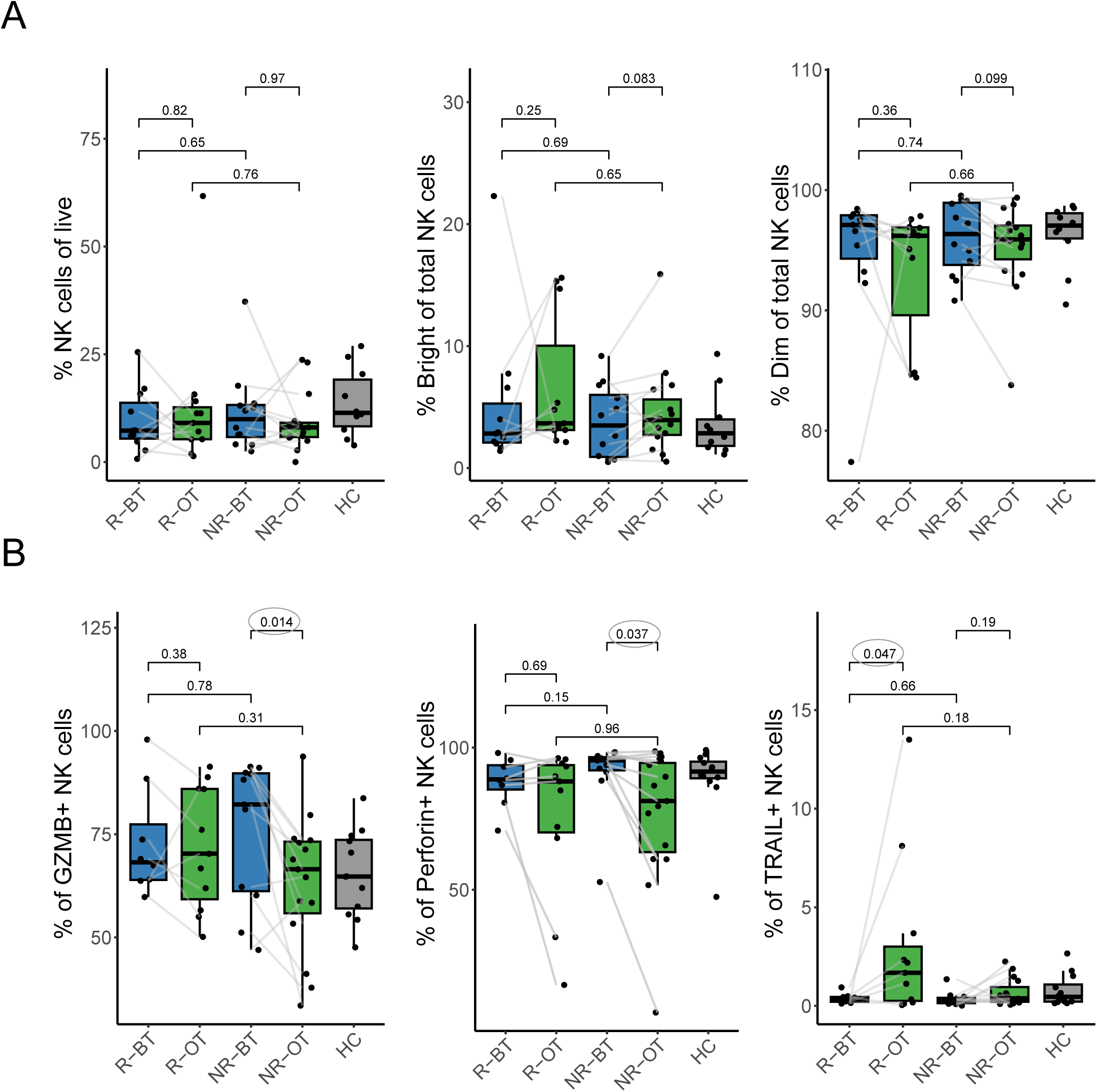
Peripheral blood NK cells phenotyping. **A,** FACS analysis revealed the percentages of living PBNK (CD3- CD14- CD19- CD123- CD56+) cells from total living PBMCs (left), CD56dim PBNK cells (CD3- CD14- CD19- CD123- CD56dim CD16+) from total PBNK cells (middle) and CD56bright PBNK cells (CD3- CD14- CD19- CD123- CD56bright CD16-) from total PBNK cells (right) in BT-NR, BT-R, OT- R and OT-NR patients. **B,** FACS analysis revealed the percentages of Granzyme B+ PBNK cells (left), Perforin+ PBNK cells (middle) and TRAIL+ PBNK cells (right) in BT-NR, BT-R, OT-R and OT-NR patients. Two-sided Wilcoxon test, circle = p<0.05 BT, Before Treatment; OT, On Treatment; NR, non-responder; R, responder

### NK cells are peripheral to the tumour bed and confined to the stroma in NR lesions

The association between cytotoxic NK cell abundance and lack of response to ICB in multiple independent cohorts is in apparent contradiction with a previous studies linking NK cells to response to anti-PD-1 therapy^14^.

Because the proximity between effector immune cells and their targets strongly correlates with response^41^, we hypothesized that differential spatial distribution of NK cells between Rs and NRs may explain this conundrum. The distribution pattern of lymphocytes within each of MM lesions was classified as “absent” (cold), “non-brisk” (immune-excluded) and “brisk” (hot)^42^ by a pathologist. As expected, we observed that the abundance of the various CD8+ T subsets (from scRNA-seq data) was significantly associated with a “brisk” infiltration pattern. Surprisingly, NK cells were most numerous in biopsies exhibiting a “non-brisk” phenotype (**Figure 4A**). We did not observe any significant associations between the remaining immune cell types and the immune infiltration pattern (**Figure S4A**). Therefore, these data indicate a potential role for NK cells in shaping the lymphocyte infiltration pattern in melanoma lesions.

**Figure 4:**
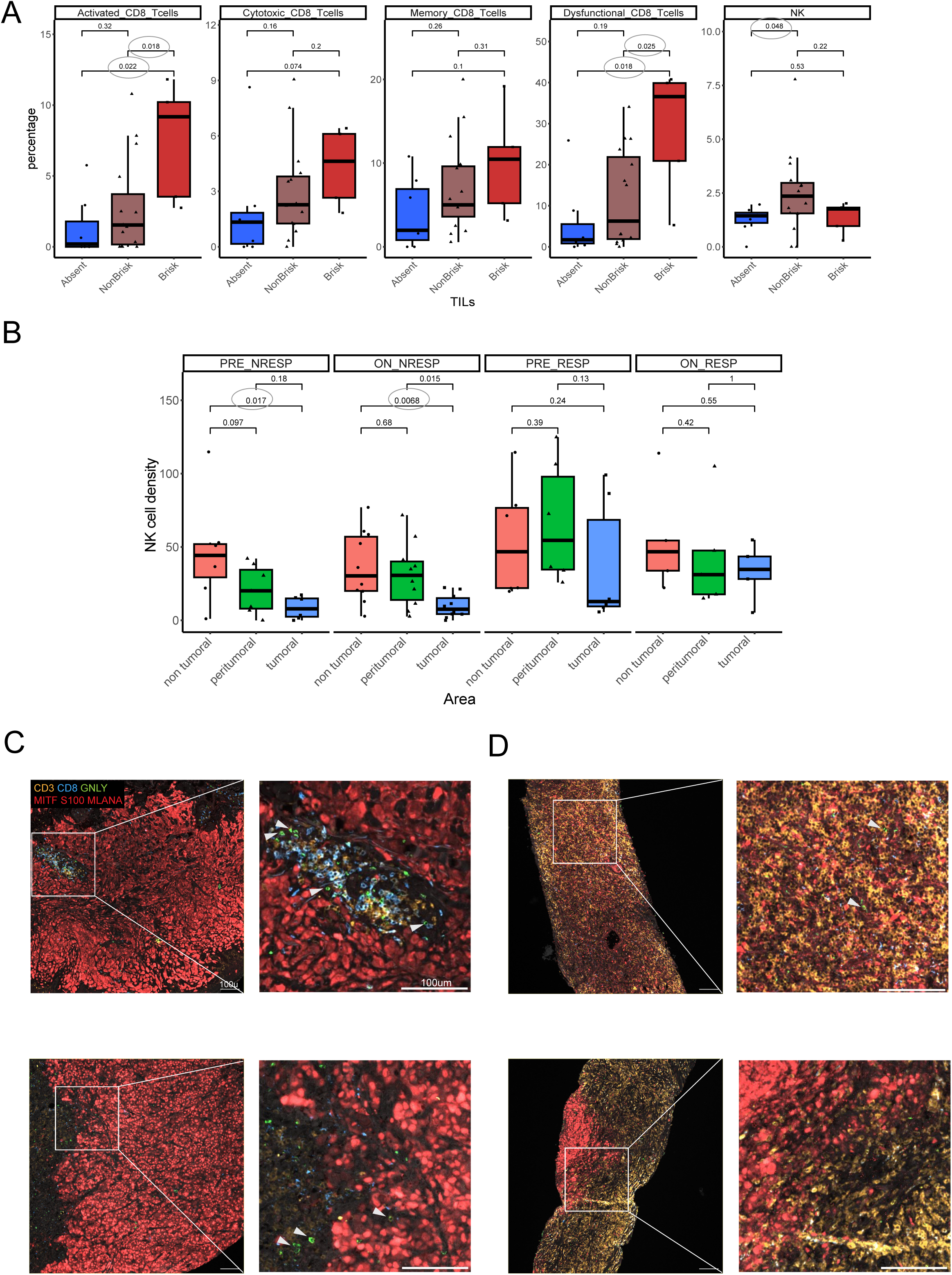
Spatial mapping of NK cells. **A,** Percentage of CD8 T-cell subsets and NK cells out of all immune cells plotted among three tumour infiltrating lymphocytes (TILS): “absent” (cold), “non-brisk” (excluded) and “brisk” (hot), two-sided Wilcoxon test, circle = p<0.05. **B,** Density per square milimiter (sqmm) of NK cells compared between three different tumoural regions, response and timepoint, two-sided Wilcoxon test, circle = p<0.05. **C,** Example of two lesions from a Non-Responder where NK cells were detected in the immune infiltrating patches (top) and at the rim of the rumour (bottom). **D,** Example of two lesions from a Responder where NK cells were intermingled with other immune cells (top) or not detected at all (bottom). Green = GNLY, red = melanoma, orange = CD3, blue = CD8 and arrows point NK cells.

To gain further insight into the spatial distribution of the NK cells, we performed high multiplexed immunofluorescence (mIF) in 17 NR samples (BT, n=7; OT, n=10) and 11 R samples (BT, n=6; OT, n=5), phenotyping individual cells with up to 41 immune, melanoma and stromal markers. Guided by our scRNA-seq data, which showed that GNLY was only expressed in NK and Cytotoxic CD8Tcells, NK cells were identified by mIF as CD45+ CD3- and GNLY+ cells (**Figure 1C**).

Interestingly, we found that NK cells display a clearly distinct spatial distribution in R and NR lesions. Each lesion was divided into tumoral, peritumoral and non-tumoral areas, as described previously^43^. NK cells were mainly localized in the non-tumoral and peri-tumoral areas in NR lesions and this differential spatial distribution was even more pronounced in the OT samples. In contrast, NK cells were evenly distributed across the three areas in Rs and this across both timepoints (**Figure 4B**). These data therefore suggest that NK cells accumulate outside the tumoral area in NR patient melanoma. Their increased abundance in non-brisk samples indicated that they are particularly enriched in lesions harbouring an “immune-excluded” phenotype. Visual examination further confirmed that NK cells preferentially colocalized with CD8+ T cells at the tumour margin in NR lesions (**Figure 4C**) In Rs, NK cells were part of a dense immune infiltrate which surrounded and interspersed with melanoma cells (**Figure 4D, top panel**) or were absent (**Figure 4D, bottom panel**).

### NK cell ablation increases ICB response in a mouse model exhibiting an immune-excluded phenotype

NK are key effector cells of innate immunity and were shown to play an important role in mediating response to anti-PD-1 therapy, including in a subset of tumours downregulating MHC class I expression^14^ (ref). However, given our findings that these cells are enriched at the tumour margin in lesions from NRs, we hypothesized that these cells may contribute to ICB resistance in lesions harbouring an immune excluded phenotype. Performing head-to-head comparison of the immune cell topography of various allograft mouse models of melanoma we identified a model exhibiting an immune excluded phenotype i.e., NRAS^Q61K/°^;Ink4a^−/− 44^(**Figure 5A**). Notably, we had previously shown that this model, which is driven by mutant NRAS^Q61K^ mutation, also harbours a cellular architecture that closely resembles what is observed in human biopsies^44^. In this model, immune (CD45+), CD3-positive T cells and NK cells (NCR1+) are confined to the peritumoral area in the tumour stroma and do not penetrate the parenchyma of the tumours (**Figure 5B**). Administration of anti-PD1 had only a marginal impact on the growth of these tumours as well as on progression free survival (**Figure 5C-D**). Likewise, systemic depletion of NK cells with the depleting antibody targeting aNK1.1 did not to impact neither tumour growth nor progression free survival (**Figure 5C-D**). Flow cytometry using both NK1.1 and Ncr1 confirmed depletion of NK cells in the blood, spleen and tumour (**Figure S5A**). Strikingly, combining NK cell depletion with anti-PD1 completely suppressed tumour growth and extended progression free survival. In sharp contrast, NK cell depletion in an allograft mouse model which harbours an immune infiltrated phenotype (**Figure S5B-C**), such as YUMM5.2^45^, decreased the sensitivity to ICB (**Figure 5E-F**). Similar results were obtained in yet another immune infiltrated allograft model, YUMM1.7 (data not shown). Note that NK cell depletion alone did not alter tumour growth, nor progression free survival, in these models (**Figure 5C-F**). These data indicated that, whereas NK cells contribute to the efficacy of ICB in melanoma lesions presenting an immune infiltrated phenotype, these cells can actively contribute to ICB resistance in melanomas harbouring an “immune excluded” phenotype.

**Figure 5:**
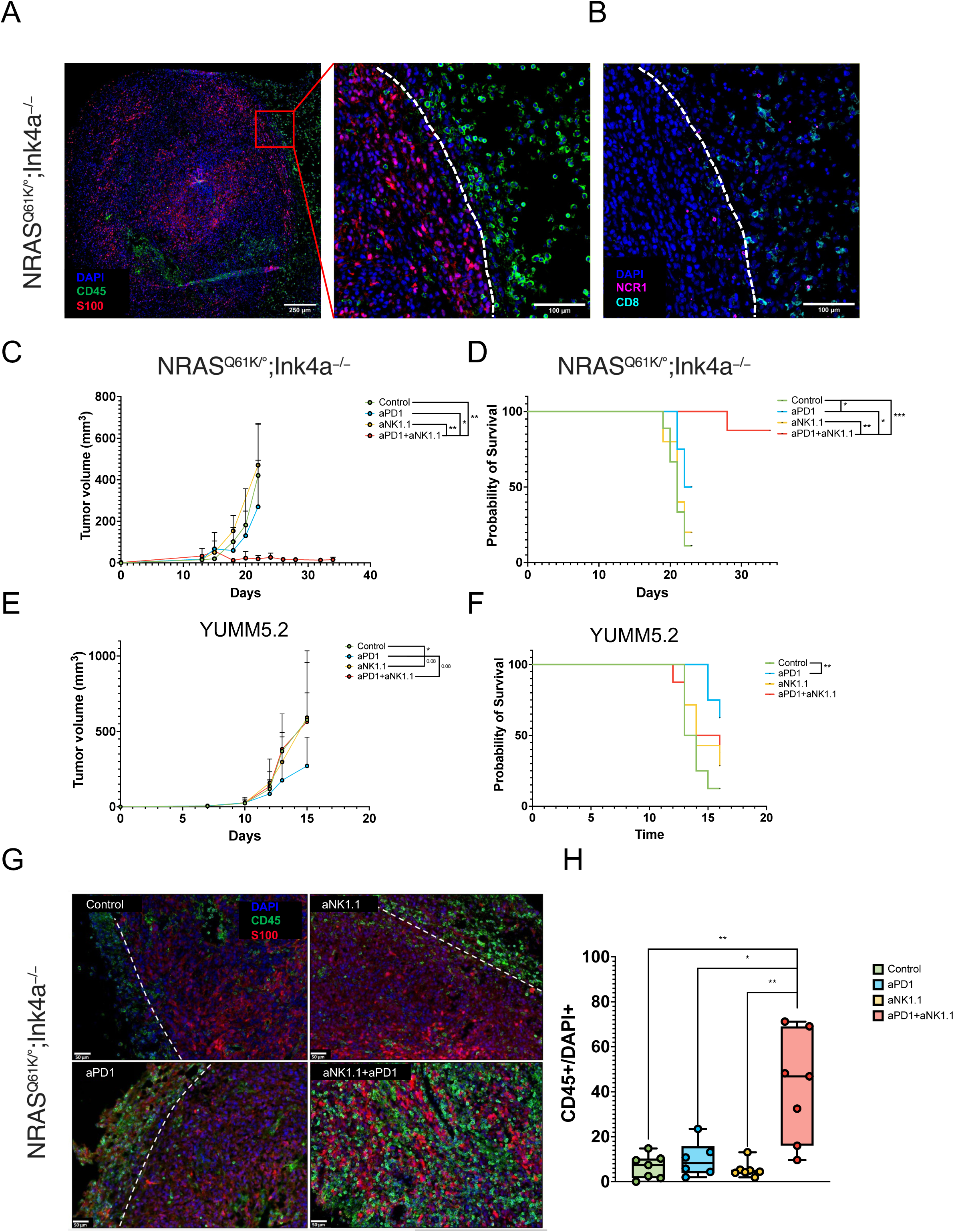
NK cells as modulators of ICB sensitivity in mice. **A,** Representative immunofluorescence image of NRAS^Q61K/°^;Ink4a^−/−^ tumour in control conditions (left panel). Melanoma cells are marked by S100 positivity, while immune cells are marked by CD45 positivity. A zoom on the tumour/immune rim is presented in the right panel. **B,** Immunofluorescence image of the same region depicted in (A), where NCR1 positivity marks NK cells and CD8 positivity marks CD8 T cells. **C,** Growth curve of NRAS^Q61K/°^;Ink4a^−/−^ tumours in C57bl/6 female mice. Data are represented as tumour size mean with corresponding standard deviation (Welch-corrected two sided t-test, * p < 0.05, ** p < 0.01; control n = 10; aPD1 n = 8; aNK1.1 n = 5; aPD1+aNK1.1 n = 9). **D,** Kaplan Meier curve of NRAS^Q61K/°^;Ink4a^−/−^ tumour progression-free survival (i.e., the time needed for the tumour to reach 300mm^3^). **E,** Growth curve of YUMM5.2 tumours in C57bl/6 male mice. Data are represented as tumour size mean with corresponding standard deviation (Welch-corrected two sided t-test, * p < 0.05; control n = 8; aPD1 n = 8; aNK1.1 n = 7; aPD1+aNK1.1 n = 8). **F,** Kaplan Meier curve of YUMM5.2 tumour progression-free survival (i.e., the time needed for the tumour to reach 300mm^3^). **G,** Representative immunofluorescence image of NRAS^Q61K/°^;Ink4a^−/−^ tumours growing across the different cohorts. The white dashed line indicates the rim between tumour and immune cells, which disappears upon NK cell depletion. **H,** Immunofluorescence-based quantification of the immune infiltrate in NRAS^Q61K/°^;Ink4a^−/−^ tumours growing across the different cohorts (Welch-corrected two sided t-test, * p < 0.05, ** p < 0.01; control n = 7; aPD1 n = 6; aNK1.1 n = 7; aPD1+aNK1.1 n = 7).

Immunostaining and histopathological analyses revealed that treatment with either anti-NK1.1 or anti-PD1 alone did not affect the distribution pattern of immune cells in the NRAS* mutant immune-excluded model (**Figure 5G** **and Figure S5D**). The combination of these antibodies, however, resulted in a massive increase in immune infiltration (**Figure 5G-H**). In fact, this effect was so dramatic that in most samples there was hardly any viable tumour cells left (**Figure S5E).** Immune fluorescent staining and flow cytometry indicated that the immune infiltrate consisted mainly of (Pd1+) cytotoxic T lymphocytes (data not shown). Moreover, a sharp increase in Pd1+ Cd8+ T cells was observed in the blood of tumor-bearing mice treated with the combination of aPD1/aNK1.1 (data not shown), indicating that the effect of NK cell depletion in combination with ICB results in systemic (re)circulation of cytotoxic T cells. Together, these data highlight contrasting roles of NK cells in modulating response to ICB depending on the immune cell topography and establish an unexpected causative link between accumulation of NK cells at the peritumoral area and intrinsic resistance to ICB.

## Discussion

With the enclosed study we provide a comprehensive characterization of the immune compartment in treatment naïve human melanomas and its evolution after a short exposure to ICB. We identified >20 melanoma-associated immune cell subsets, including various CD8+ T cell, monocyte, macrophage and dendritic cell subclusters. This dataset constitutes a very valuable resource for the entire melanoma community and beyond.

Consistent with previous studies^6,7,29^ we observed a higher proportion of various subsets of CD8+ T cells in lesions from patients benefiting from this treatment. We confirmed the association of Activated, Dysfunctional and Memory CD8+ T cells with ICB response in BT biopsies^6^.

Subclustering the macrophage population allowed us to discern several previously described (tumour-associated) macrophage subpopulations, including a cell state characterized by higher levels of *C1Q* and *SPP1*^30,46^. A C1Q+ population has been associated with poor prognosis in various cancer types and proposed to promote an immunosuppressive microenvironment^47^. Likewise, the presence of SPP1+ macrophages has also been linked to poor prognosis and resistance to ICB. This cell population seemingly resides in hypoxic areas where it interacts with cancer-associated fibroblasts resulting in remodelling of the extracellular matrix^31,48,49^. There was, however, no significant association between these macrophage subpopulations and therapy outcome in our cohort. This could be due to differences in the design of the respective studies, and/or in the size of the cohorts analysed.

Unique to our study was a significant increase in NK cells in on-treatment lesions from patients who failed to respond to ICB. Importantly, we could validate this finding in independent melanoma cohorts and, even in a breast cancer cohort indicating that this observation may be generalizable to other cancer types. Several of the most discriminative genes (e.g., *GNLY*, *PRF1*, *GZMB*) of the NK cell cluster indicated that these cells may exhibit a cytotoxic phenotype. Moreover, further classification of this cluster into NK_CYTO_ and NK_REST_, as previously described in a breast cancer cohort^39^ indicated that the association between NK cell abundance and lack of response is driven by the NK_CYTO_ phenotype, characterized by *FCGR3A* (CD16), *FGFB2* and cytotoxic molecules (*GNLY*, *PRF1*, *GZMB*). This unexpected observation starkly contrasts with previous reports in which NK cell abundance was linked to response to aPD1 therapy^14^, and another study showing that expression of NK cell-related genes is predictive for ICB response and prognostic for survival in MM patients^15^. A rational explanation for these apparently conflicting results was provided by our spatial proteomics analysis, which indicated that the increase in NK cell abundance was confined to regions located outside of the tumour bed. This increase in NK cell abundance in the non-tumoral area of NRs OT was not mirrored in the peripheral blood. Multiparametric flow cytometry indeed failed to highlight a significant difference in the proportion of total or CD56^dim^ NK cells in the blood of the R versus NR cohort. This observation agrees with a previous report showing that low baseline level of total or CD56^dim^ PBNKs were associated with response to aCTLA4 but not aPD1 therapy^50^. Similarly, to a previously published integrated scRNA-seq analysis, extensive immunophenotyping did not reveal the presence of heterogeneous NK cell populations in the peripheral blood of our patients, nor significant differences across ICB outcomes or sampling time points^51^. We did observe, however, a decrease in cytotoxic NKs in the blood of NRs early during treatment, indicating a possible recruitment of these cells into peripheral tissues.

A robust biomarker that reliably predicts response to ICB in MM patients is still lacking. The dynamic increase of cytotoxic NK cells at the tumour border of NR lesions at an early on-treatment time point maybe exploited for the development of a multiparametric immunostaining-based predictive biomarker. Given our observation that this increase may be occurring in other tumour types, such as breast cancer may even further increase the potential of such a composite biomarker.

This increase in NK cells may also highlight an active role in promoting intrinsic resistance to ICB. Consistent with this possibility, we show that depletion of NK cells in a mouse model that exhibit an immune-excluded, but not immune-infiltrated, phenotype drastically increases response to ICB. Future studies aiming at dissecting the molecular mechanisms underlying this unexpected role of NK cells may help develop innovative therapeutic approaches to overcome resistance to ICB in MM and beyond. Moreover, these results also indicate that therapeutic strategies employing NK cells to overcome ICB limitations, such as CAR-NK cell therapy, need to be considered with caution and take the immune cell topography of the target lesions into account.

## Methods

### Patient samples

Tumour biopsies were collected as part of a non-interventional prospective study investigating transcriptomic changes upon immune checkpoint inhibition (Prospective Serial biopsy collection before and during immune-checkpoint inhibitor therapy in patients with malignant melanoma; SPECIAL). Patient recruitment and sample collection is previously described^23^. In brief metastatic lesions of treatment-naïve patients with advanced (st. III and st. IV) cutaneous melanoma, treated with anti-PD-1 based ICB (either in (neo-)adjuvant or inoperable setting) were biopsied right before the administration of the first and second ICB cycle. Demographic, clinical, histopathological and genetic information was collected at baseline. Patients with unresectable disease were stratified as responders (complete remission, partial remission) and non-responders (stable disease, progressive disease) based on RECISTv1.1. best overall response, whereas patients treated with curative intent were stratified according to pathological response assessment at tumour resection. Written informed consent was obtained from all patients. All study procedures were in accordance with the principles of the Declaration of Helsinki, applicable Belgian law and regulations, and approved by the UZ Leuven Medical Ethical Committee (S62275 and S62927).

### Single-cell RNA-sequencing oh human samples

Methods for tumour dissociation, library construction and sequencing, scRNA-seq data acquisition and analysis were described previously^14^.

Immune cells were identified using a gene set score of >0.1 for the Immune score from Jerby-Arnon et. al.^15^, in addition to an inferred mean CNV score < 0.1. Next, the SCTransform was applied regressing out mitochondrial read percentage and cell cycle scores and the data integration (by samples) was performed using R package Harmony v. 1.0^30^. The main immune cell types (n = 17) were identified after first round of unsupervised Louvain clustering and marker gene calling. Next, all CD8+ T cell clusters were then reclustered together for deeper characterisation. Myeloid and Plasmacytoid dendritic cell clusters were further reclustered, followed by reclustering of Macrophages separately and Dendritic cells together with Monocytes. The marker genes of each identified cluster were calculated using FindAllMarkers function in Seurat (Wilcoxon Rank Sum test).

Stromal component (Cancer Associated Fibroblasts (CAFs) and Endothelial Cells (ECs)) were identified based on gene signatures from Jerby-Arnon et. al.^15^)

### Validation of NK cells in independent single cell RNA sequencing dataset

The Transcript Per Million (TPM) normalized Sade-Feldman et al. dataset was downloaded from the GEO portal (accession number GSE120575)^6^. Importantly, samples in this study were recruited from a mixed cohort which included patients treated anti-CTLA-4 in monotherapy. Therefore, we restricted our validation analyses to patients treated with only aPD-1 and in combination with anti-CTLA4 treatment. Cells with > 1000 & < 7500 expressed genes were selected for further analysis. NK cells in this independent dataset were identified within G8 (Cytotoxicity) cluster based on an AUCell score of the differentially expressed genes in the NK cell cluster in our dataset, compared to the CD8Tcells (Table S3). Additionally, cells positive for CD8A were excluded. The cell percentage positive for this score was compared between responders and non-responders using Wilcoxon Rank Sum test.

CIBERSORTx was used to infer main TME components: B cells, Endothelial Cells (EC), Macrophages, CD8+ T cells, T regulatory cells (Tregs), Malignant cells, Cancer Associated Fibroblasts (CAFs), Dendritic Cells (DCs), CD4+ T cells, and Natural Killer cells from the Riaz, et al. bulk RNA sequencing dataset (accession number GSE91061)^37^. Count matrices were subsampled for 15.000 cells and used to generate cell type expression signatures with default setting except for the minimal expression parameter, which was set to 0. The TME cell types were imputed with default parameters with batch correction mode.

CIBERSORTx was run with rmbatchSmode = TRUE, nsampling = 30, QN= FALSE.

To score the NK cells signatures from NK cells identified in the breast cancer cohort we run the differential gene expression of all immune cells (method), compared the top 50 genes from NK_CYTO_ and NK_REST_ and selected only unique gene sets per cell type. These gene sets were further used to calculate AUCell scores. The AUCell scores were further used to calculate the percentage of “NK cells high” in NK cyto and rest signatures.

### Flow cytometry of human PBMCs

Patient blood samples were obtained at the time of tumour biopsy, in 3x 10 mL EDTA tubes: 28 NR samples (BT, n = 13; OT, n = 15) and 23 R samples (BT, n = 12 OT, n = 11). In addition, we collected blood from age- and sex-matched controls (n = 10; Table S1B). PMBCs were isolated using Ficoll-Paque density gradient centrifugation. First, blood was diluted 1:1 in sterile Dulbecco’s phosphate-buffered saline (DPBS; Thermo Fischer Scientific Cat #14-200-075). This mixture was carefully layered on top of 10 mL of sterile Ficoll-Paque Plus (Sigma-Aldrich, Cat# GE17-1440-02) and centrifuged 30 min. At 400G at room temperature, after which the diluted plasma layer on top was pipetted off and the interface layer containing the PBMCs was carefully collected. Cells were resuspended in DPBS and washed twice before cryopreservation in 10% dimethyl sulfoxide (DMSO; Sigma-Aldrich Cat# 472301) in fetal bovine serum (FBS, VWR, Cat # S1810-500) at −80°C.

For flow cytometry, cells were thawed (0.5×10^6^ per staining), incubated with human FcR-block (5µl/sample, Miltenyi Biotec) and stained with monoclonal antibodies (list of human antibodies is shown in Table S8). Dead cells were excluded using Zombie Aqua 516 (1:1000, Biolegend) or Fixable Viability Stain 620 (1:25000, BD biosciences) Flow cytometric analysis was performed on a BD LSR Fortessa X20 with DIVA software. Results were analysed with FlowJo (LLC, V10).

### Multiplex imaging and digital pathology

Multiplex Iterative Labeling by Antibody Neodeposition (MILAN) platforms were carried out as previously described^23^ on formalin fixed paraffin embedded tissue biopsies: 17 NR samples (BT, n = 7; OT, n = 10) and 11 R samples (BT, n = 6 OT, n = 5), MILAN entails multiple rounds of indirect immunofluorescent staining using unconjugated antibodies, followed by imaging and antibody removal via detergent and a reducing agent. Image analysis was performed as previously described^42^. Briefly, stains were visually evaluated for quality by an experienced pathologist. Flat field correction was performed using a custom implementation of a publicly described methodology^52^. Consecutive staining rounds were registered using a publicly available algorithm^53^. Tissue autofluorescence was subtracted using a baseline image stained only with a secondary antibody. Nuclear cell segmentation was performed using a fine-tuned version of StarDist^54^ Springer International Publishing.). Phenotypic identification was done based on prior knowledge using a two-tiered approach combining different clustering methods with functional analysis. Finally, the tissue was digitally reconstructed to perform spatial analysis. Antibodies used in this experiment are listed in Table S8. NK cells defined as CD3-GNLY + cells were counted per square milimiter (sqmm) unless otherwise specified.

### Mouse experiments

Male and female C57BL/6 were purchased from KU Leuven Mouse Facility and housed at 22°C ± 2°C, 55% ± 10% relative humidity, and with 12 hours d/light cycles in mouse facilities at the KU Leuven Mouse Facility (Leuven, Belgium). *In vivo* studies were performed after approval from the KU Leuven Ethical Committee for Animal Experimentation (ECD number 188/2022).

2.5*10^5^ NRAS;Ink4a cells were subcutaneously injected into 8-12 weeks old female C57BL/6 mice, while 2.5*10^5^ YUMM5.2 cells were subcutaneously injected into 8-12 weeks old male C57BL/6 mice. Tumour volume was monitored every other day and calculated using the formula V = (width * length^2^)/2. As soon as tumours reached ∼20mm^3^ (indicatively corresponding to 3mm x3mm mass), mice were administered either anti-PD1 (clone RMP1-14) or IgG2a control isotype (BioXCell; 200μg in 200μl, twice per week). To perform NK cell depletion, mice were administered anti-NK1.1 depleting antibody (clone PK136) 3 days before tumour injection and every five days after tumour injection. Mice were sacrificed (by cervical dislocation) in all the cohorts once the tumour volume in control cohort reached ∼500mm^3^ volume. We decided to maintain longer uniquely the anti-PD1^+^ anti-NK1.1 cohorts in NRAs;Ink4a model, to appreciate the extent of prognosis improvement upon treatment. Progression-free survival was computed as the amount of time needed for the tumour volume to exceed 300mm^3^.

#### Immunofluorescence on cryosections and FFPE sections

NRAS^Q61K/°^;Ink4a^−/−^ tumors were resected, rinsed in ice cold DPBS, fixed in 4% formaldehyde for 2 hours, washed in DPBS and placed in 30% sucrose in DPBS overnight. Next, tissues were embedded in Tissue-Tek® O.C.T. Compound (Sakura® Finetek, #4583) and stored at −80°C. Sections of 10 μm were cut using the Thermo Scientific CryoStar NX70 Cryostat. For immunofluorescence, tissue sections were air dried, rinsed in DPBS and fixed for 10 min in 4% paraformaldehyde. They were washed in DPBS for 5 min, permeabilized in 1% Triton X-100 in DPBS for 10 min and washed in DPBS for 5 min again. Next, sections were incubated in blocking buffer (1% BSA, 10% donkey serum, 0.1% Triton X-100 in DPBS) for 45 min. All these steps were carried out at RT. Primary antibodies against CD3 (Dako cat. no A0452; 1: 200), CD8 (Novus Biologicals cat. no NB200-578; 1:200), NKp46 (R&D Systems cat. no AF2225; 1:200), CD45 (Novus Biologicals cat. no NB100-77417SS; 1:400) and S100 (Dako cat. no Z0311; 1:200) were incubated overnight at 4°C in antibody diluent (1% BSA, 0.1% Triton-X DPBS). Sections were washed in DPBS for 10 min and incubated with secondary antibodies (Donkey anti-rabbit Life Technologies cat. no A31573, 1:400; Donkey anti-rat ThermoFisher Scientific cat. no A21208, 1:400; Donkey anti-goat ThermoFisher Scientific cat. no A11058, 1:400) in antibody diluent for 45 min at RT. Sections were twice in DPBS for 7.5 min at RT. Finally, nuclei were stained with DAPI (4’,6-diamino- 2-phenylindole, dihydrochloride, Thermofisher, #D3571) solution (0.5 mg/ml), diluted 1:1000 in DPBS for 5min and mounted in ProLong™ Diamond Antifade Mountant (Thermofisher Scientific, #P36961)

YUMM5.2 tumours were collected upon mouse sacrifice and immediately placed in 4% formaldehyde solution overnight. Afterwards, samples were moved to 70% ethanol and subsequently included in paraffin. Five µm FFPE tissue sections of selected samples were cut and mounted on coverslips. Deparaffinization and rehydration was performed with Leica Autostainer XL (ST5010). Afterwards, samples were fixed for 10’, and washed in TBST (3’, 3 times). Antigen retrieval was performed with Antigen Retrieval Buffer (100X Tris-EDTA Buffer, pH 9.0; Abcam) for 20’ in pressure cooker. Afterwards, samples were cooled down gradually at room temperature, washed, and blocking was performed with 1% BSA in TBST. Samples were then incubated overnight in blocking solution with antibodies recognizing CD45 (Cell Signaling cat. no 70257S; 1:200) or CD3 (Abcam cat. no ab16669; 1:100). Donkey anti-rabbit AF647-conjugated secondary antibody was added the day after upon wash (Life Technologies cat. no A31573; 1:400). Samples were then washed, DAPI counterstained (1:1000), mounted with ProLong™ Gold Antifade Mountant solution (Invitrogen), and imaged with Vectra Polaris/PhenoImager HT (Akoya).

#### FACS

Tumour, lymph nodes and spleens were resected, rinsed in ice cold DPBS, mechanically minced into small pieces with sterile scalpels and enzymatically dissociation by incubation in 750 µL digestion buffer (2 ng/mL Collagenase P, Sigma-Aldrich #11249002001, 0.8 mg/mL DNase I, Sigma-Aldrich #11284932001 in Dulbecco’s Modified Eagle Medium, high glucose with Glutamax (DMEM, Invitrogen #61965059) ofr tumours and 300μg/ml Liberase, Sigma Aldrich, #5401127001 and 1 mg/mL DNAseI 1 mg/ml in DMEM for lymph nodes and spleens) for 15 minutes in a heather-shaker at 37°C 800 rpm. Digestion buffer was inactivated by adding 750 µL DPBS. Samples were centrifuged at 300G for 5 min. at 4°C. Supernatans was discarded, and the pellet was resuspended in 500 µL ACK red blood cell lysis buffer for 5 min. at RT (Thermo Fisher, Cat#A1049201). Next, cell were centrifuged again at 300G for 5 min. at 4°C, resuspended in 4 mL 0,04% bovine serum albumin (BSA, Sigma-Aldrich, #A9647-100G) in DPBS and filtered through a 40 μm Falcon round bottom with Cell Strainer Snap Cap (Corning, #352235). After centrifugation for at 300G for 5 min. at 4°C, cells were resuspended in FACS buffer (DPBS supplied with 2% fetal bovine serum and 2 mM EDTA), counted on a LUNA™ Automated Cell Counter, and diluted to reach an estimated concentration of 10^6 cells per mL. 500 µL of sample was transferred to a fresh 5 mL Falcon tube and incubated 1:5000 with eBioscience Fixable Viability Dye eFluor 506 for 5 min. at RT. Samples were washed in FACS buffer, incubated with staining antibodies against CD45 (Biolegend cat. no 103108; 1:300), F4/80 (ThermoFisher Scientific cat. no 45-4801-82; 1:200), CD3 (Biolegend cat. no 100221; 1:300), CD8a (Biolegend cat. no 100707; 1:300), NKp46 (Biolegend cat. no 137611; 1:50), and PD1 (Biolegend cat. no 135209; 1:100) for 20 min. at RT in the dark, washed thrice in FACS buffer. After centrifugation at 300G for 5 min. at RT, cells were resuspended in 400 µL BD Cytofix/Cytoperm solution (BD Biosciences #554714) for 20 min. at 4°C in the dark. Cells were washed twice in FACS buffer and kept at 4°C in the dark. Flow cytometry was done on a BD Fortessa X-20 within 48 hours. Analysis was performed in FlowJo.

#### Statistical analysis

For the mouse experiments, sample size is reported in the corresponding figure legend. Statistical significance was obtained comparing the tumour size at the last time point measured (Welch-corrected t-test, p-values are reported in the corresponding figure legend).

## Author contributions

J.P. analysed the scRNA-seq G.B. and E.L. and N.R performed mouse experiments. Y.v.H. performed the MILAN experiments. A.A., F.B. and Y.v.H. analysed and interpreted the MILAN data. F.B. provided pathology support. E.L. and Y.v.H. curated patient clinical data. A.d.V. perfomed human blood FACS experiments under supervision of P.M. A.B. provided NK cells analysis of the breast cancer cohort under supervision of D.L. J-C.M and J.P conceptualized and designed the research study. J-C.M, J.P., E.L. and N.R. wrote the manuscript. All authors read and edited the manuscript.

## Acknowledgments

We thank prof. Charlotte Scott (VIB-UGent Center for Inflammation Research) for valuable input on the myeloid cell annotation and Bert Malengier-Devries for initiating human PBMC experiments. J.P. received a Marie Curie Fellowship (H2020-MSCA-IF- 2019, #896897) and Stichting tegen Kanker fellowship (2023/2310), A.A. was funded by FWO Postdoctoral Fellowship Senior (12ATN24N). F.B. was funded by KULeuven (EMH-D8191-AKUL/19/30 I005920N), and FWO Fundamenteel Klinisch Mandaat (EMH-D8972-FKM/20). J-C.M received funding from the VIB Grand Challenges Program (POINTILLISM), FWO/KOTK (G0B1622N) and FWO/FRS-FNRS (EOS, # 40007513). Some figures were created with BioRender.com.

## Figure legends

**Supplemental Figure S1: Dissecting the immune compartment of the melanoma ecosystem**

**A,** Uniform Manifold Approximation and Projection (UMAP) of the common marker genes of the main immune cell types.

**B,** UMAP of the main immune cell types identified by unsupervised Louvain clustering.

**C,** UMAP of reclustered Myeloid cells together, Macrophages C1Q (**D**) and Dendritic cells with monocytes (**E**).

**F,** Percentages of identified immune cell types out of all immune cells across metastatic sites (LN:Lymph Node, Subcutaneous and Skin; two-sided Wilcoxon test, circle = p<0.05) **G,** Mosaic plot representing lack of association of the metastatic site and response to ICB.

**Supplemental Figure S2: Immune cell types association with ICB response and timepoint.**

**A,** Percentages of identified immune cell types out of all immune cells across before treatment (BT) and on treatment (OT) samples and both response groups (two-sided Wilcoxon test, circle = p<0.05).

**Supplemental Figure S3: FACS gating strategy**

Representative gating strategy of PBNKs in PBMCs is shown. PBNKs are defined as CD3- CD14- CD19- CD123- CD56+.

**Supplemental Figure S4: Association of immune cells with TILs pattern**

**A,** Percentages of identified immune cell types out of all immune out of all immune cells plotted among three tumour infiltrating lymphocytes (TILS): “absent” (cold), “non-brisk” (excluded) and “brisk” (hot). Two-sided Wilcoxon test, circle = p<0.05.

**Supplemental Figure S5: NK cell ablation and response to ICB in mice**

**A,** FACS-based quantification of NK cell depletion upon aNK1.1 administration. NK cells were defined as CD45+ CD3- NCR1+ and their number was divided by the number of CD45+ CD3- cells (Welch-corrected two sided t-test, * p < 0.05; PBMC n = 3 (aPD1) + 2 (aPD1+aNK1.1); Spleen n = 2 (aPD1) + 2 (aPD1+aNK1.1); Tumor n = 2 (aPD1) + 2 (aPD1+aNK1.1)).

**B,** Representative immunofluorescence images of YUMM5.2 tumors in control conditions. Immune cells are marked by CD45 positivity (left and middle panel), while T cells are marked by CD3 positivity (right panel).

**C,** Representative immunofluorescence image of NRAS^Q61K/°^;Ink4a^−/−^ tumour in aPD1 conditions (left panel). Melanoma cells are marked by S100 positivity, while immune cells are marked by CD45 positivity. A zoom on the tumour/immune rim is presented in the right panel.

**D,** Representative immunofluorescence images of NRAS^Q61K/°^;Ink4a^−/−^ tumours in aPD1+aNK1.1 conditions. Melanoma cells are marked by S100 positivity, while immune cells are marked by CD45 positivity.

