## Supplemental Figures for "NK cells contribute to resistance to anti-PD1 therapy in immune-excluded melanomas"

### Supplemental Figure S1

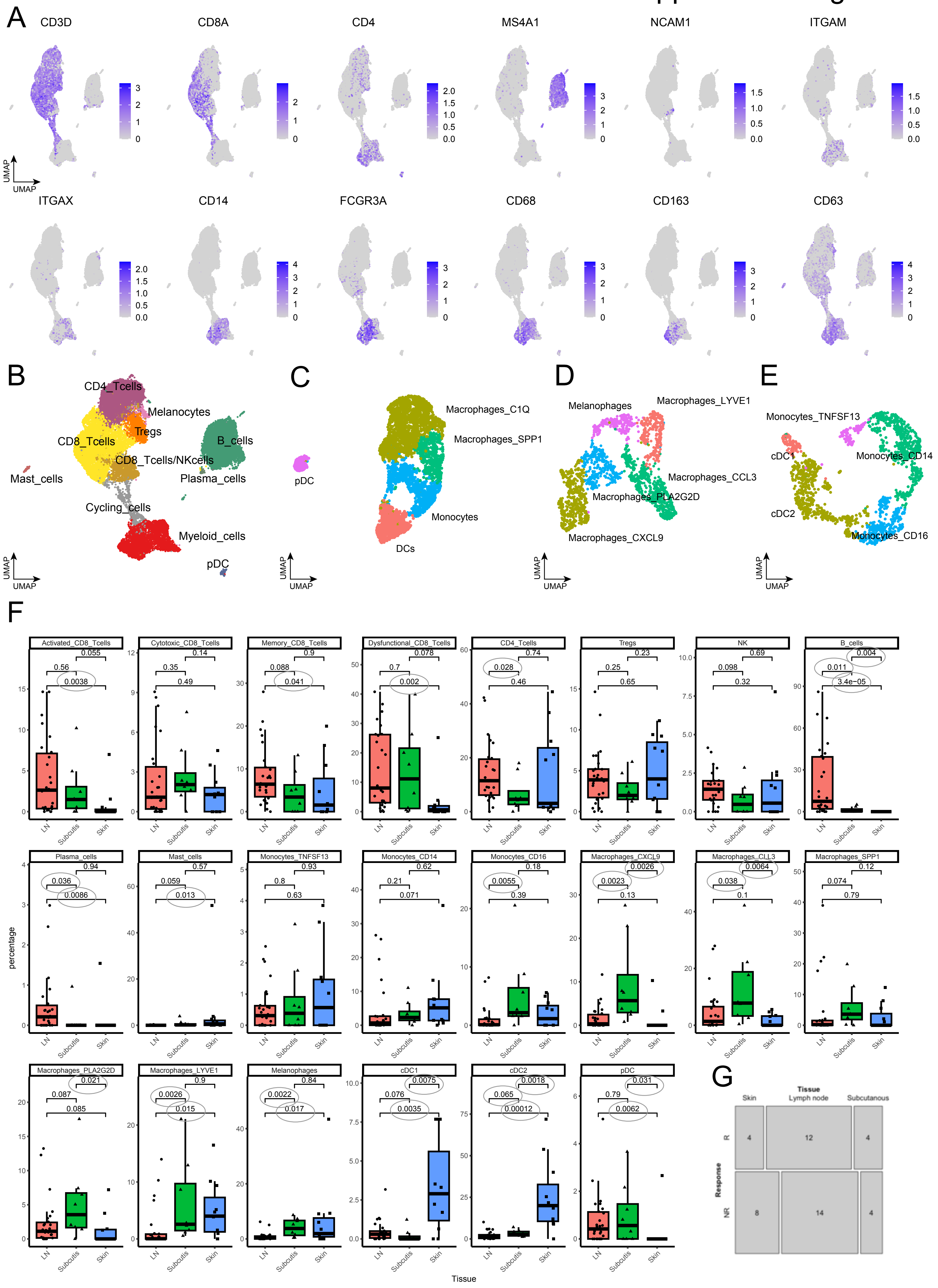

### Supplemental Figure S2

A

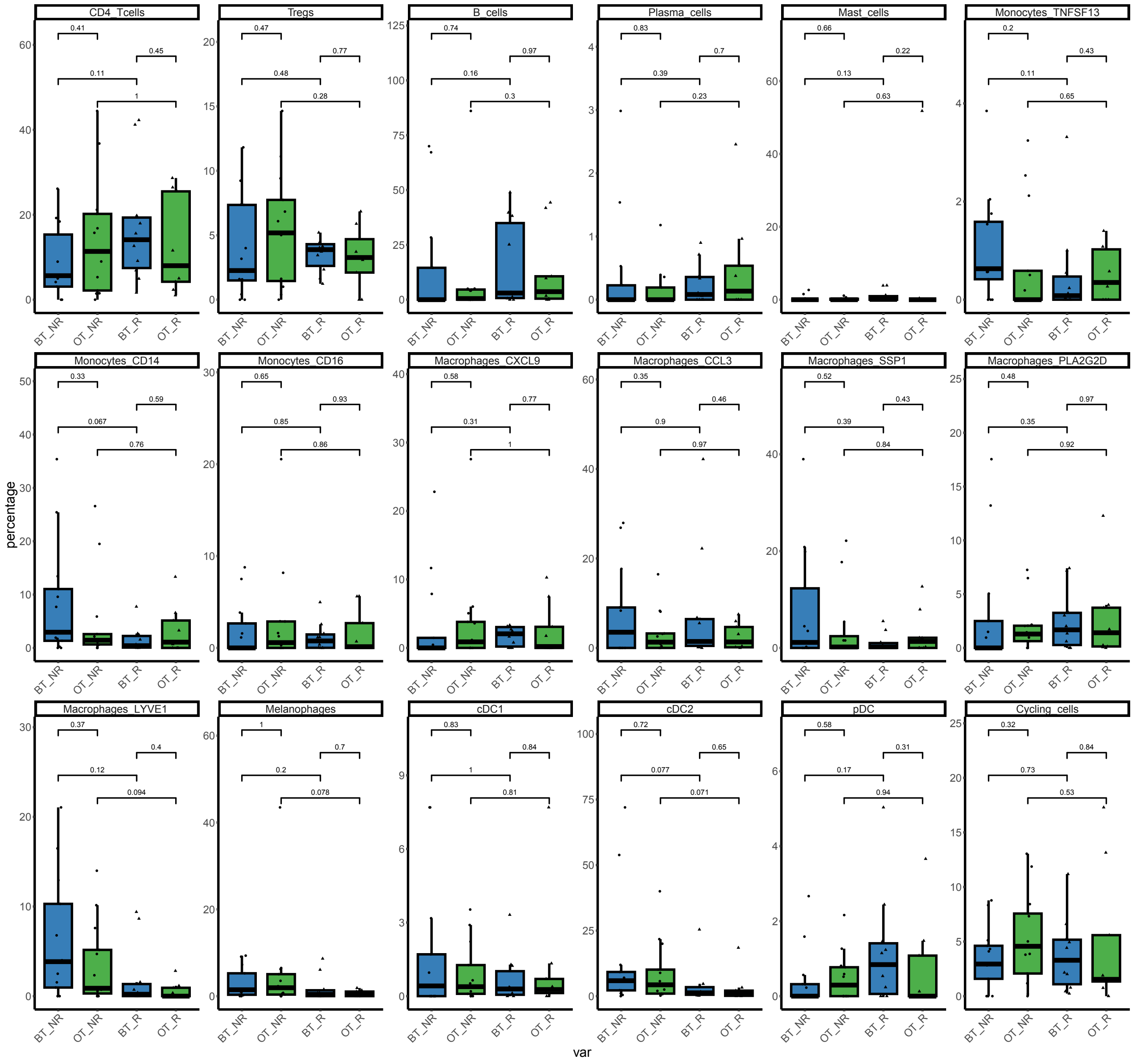

### Supplemental Figure S3

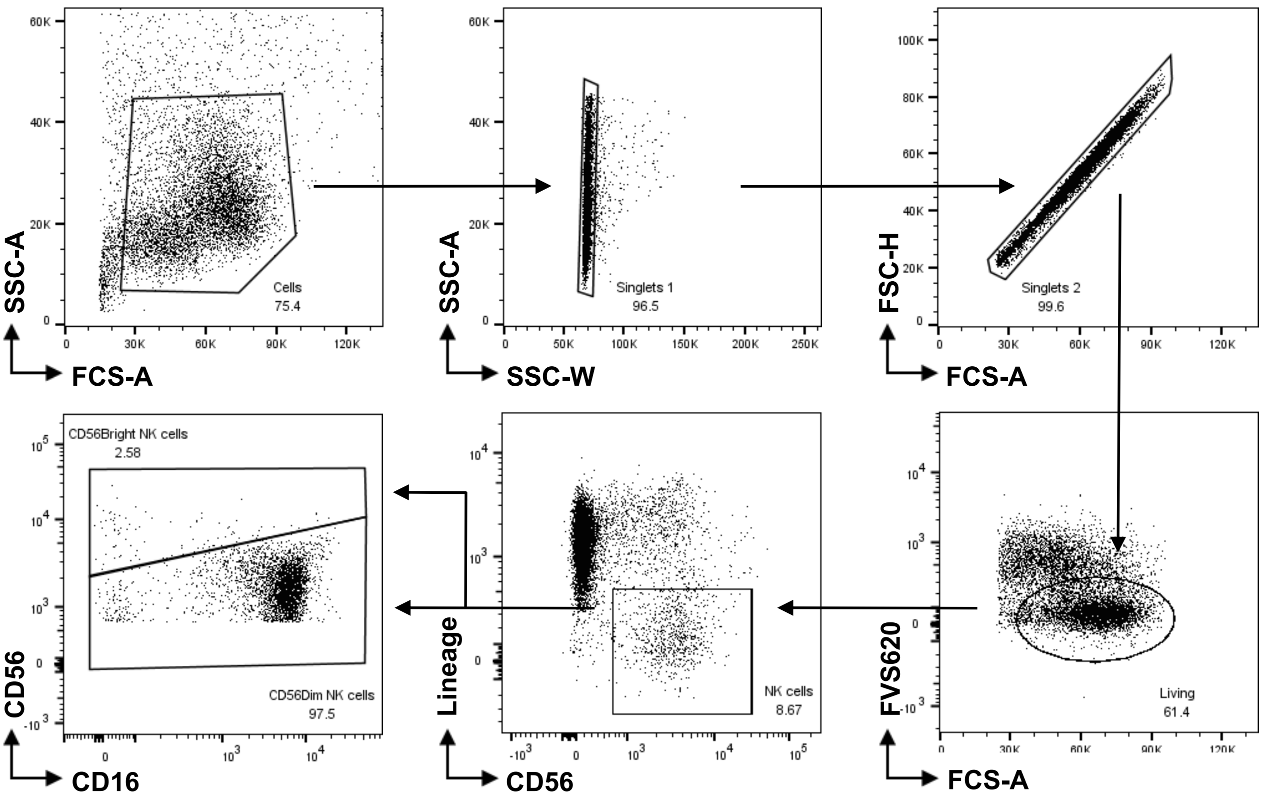

### Supplemental Figure S4

A

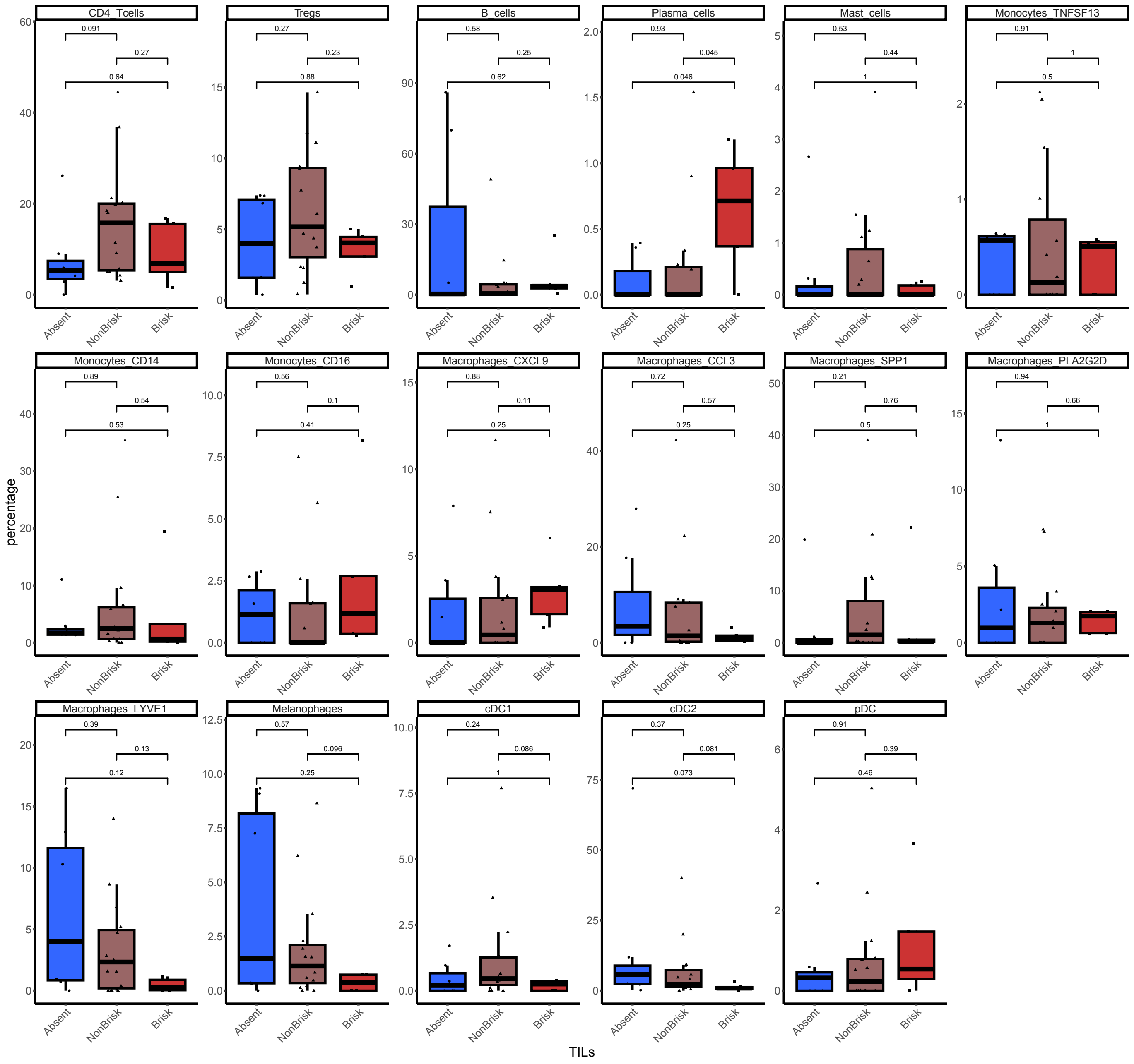

### Supplemental Figure S5

A

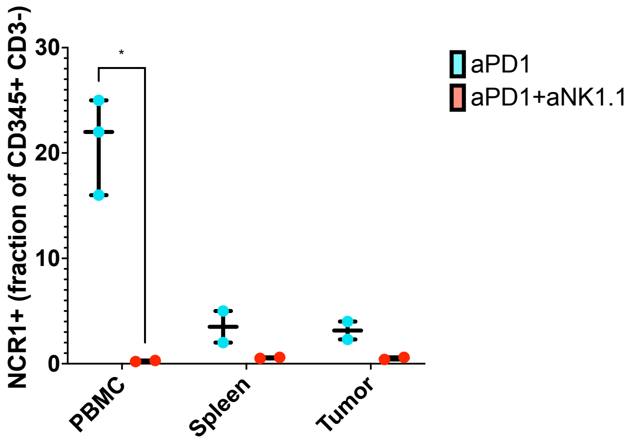


B

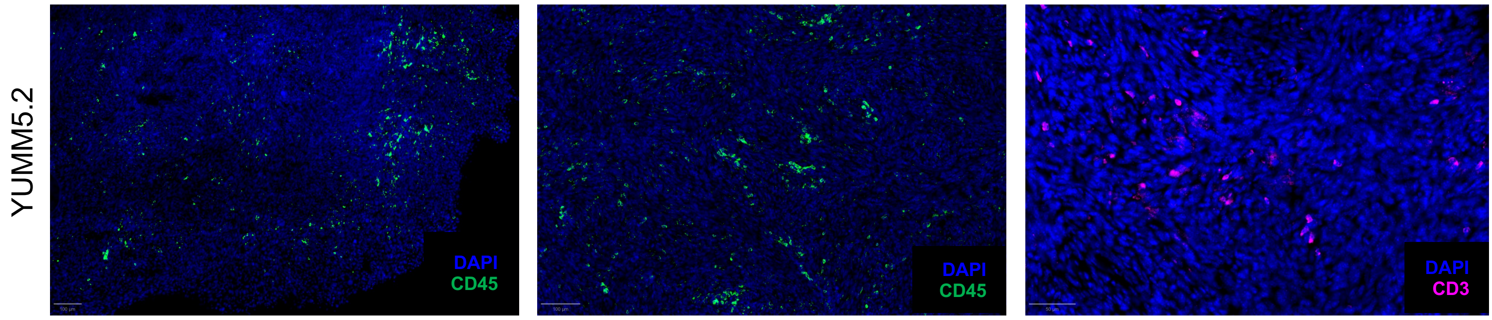


C

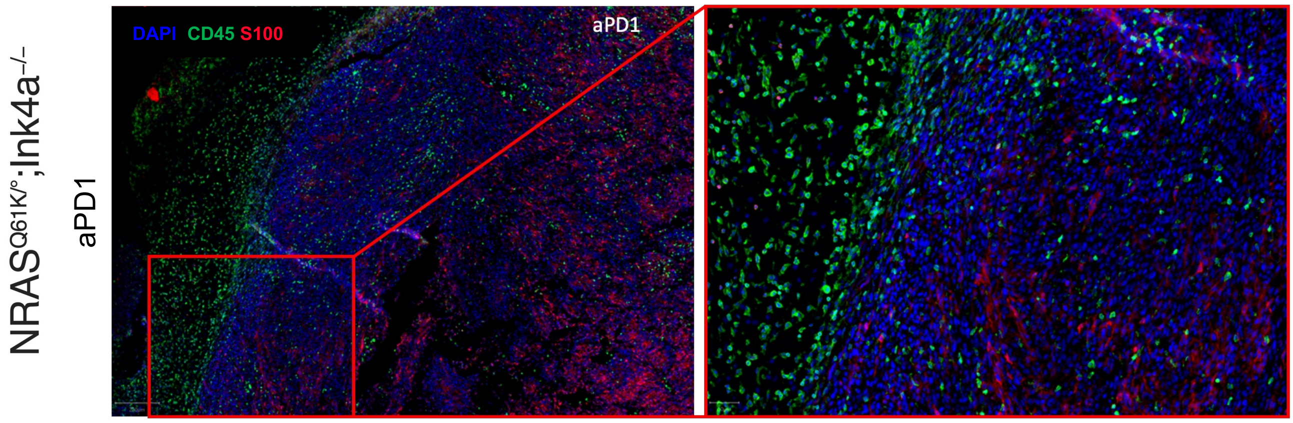


D

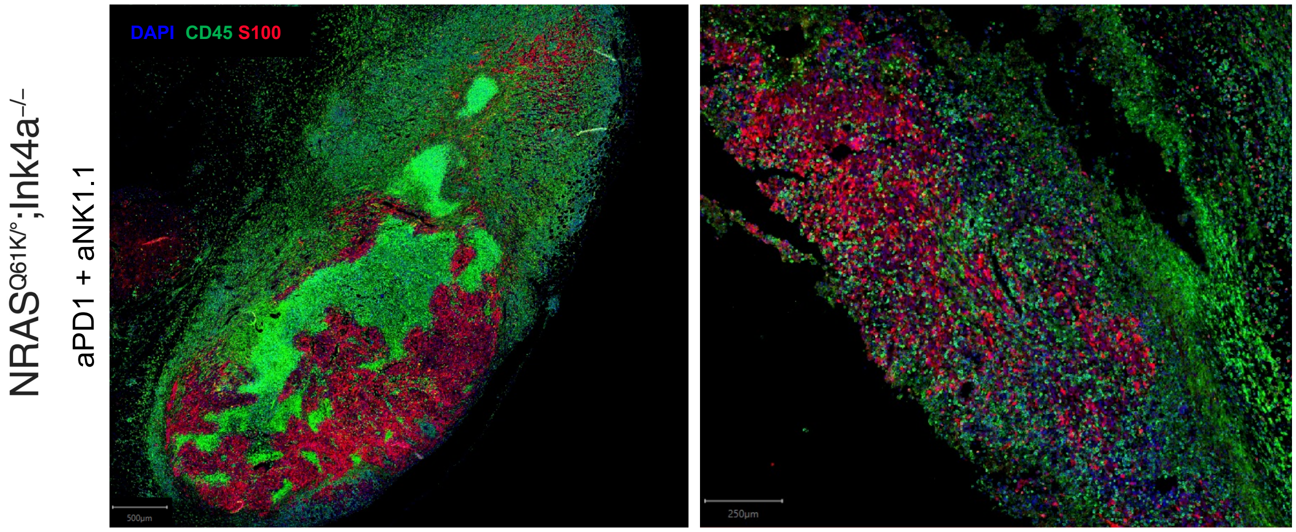
